# Converging effects of cannabis and psychosis on the dopamine system: A longitudinal neuromelanin-sensitive MRI study in cannabis use disorder and first episode schizophrenia

**DOI:** 10.1101/2024.09.12.612734

**Authors:** Jessica Ahrens, Sabrina D. Ford, Betsy Schaefer, David Reese, Ali R. Khan, Philip Tibbo, Rachel Rabin, Clifford Cassidy, Lena Palaniyappan

**Affiliations:** Integrated Program in Neuroscience, McGill University, Montreal, Quebec, Canada; Douglas Research Institute, Verdun, Quebec, Canada; Robarts Research Institute & Lawson Health Research Institute, London, Ontario, Canada; Department of Medical Biophysics, Schulich School of Medicine and Dentistry, Western University, London, Ontario, Canada; Department of Psychiatry, Dalhousie University, Halifax, Nova Scotia, Canada; Renaissance School of Medicine at Stony Brook University, Stony Brook, New York, USA

## Abstract

**Importance:** Despite evidence that individuals who use cannabis early in life are at elevated risk of psychosis and that the neurotransmitter dopamine has a role in both conditions, the mechanism linking the two conditions remains unclear.

**Objective:** To use neuromelanin-sensitive MRI (neuromelanin-MRI), a practical, proxy measure of dopamine function, to assess whether a common alteration in the dopamine system may be implicated in cannabis use and psychosis and whether this alteration can be observed in cannabis users whether or not they have a diagnosis of first-episode schizophrenia.

**Design, Setting, and Participants:** This longitudinal observational study recruited participants from 2019 to 2023 from an early intervention service for psychosis in London, Ontario, Canada. The sample consisted of 25 participants with cannabis use disorder (CUD) and 36 participants without CUD (nCUD), of which 28 had first-episode schizophrenia (FES). One-year follow-up was completed for 12 CUD and 25 nCUD participants.

**Main Outcomes and Measures:** Neuromelanin-MRI contrast within the substantia nigra (SN) and within a subregion previously linked to psychosis severity (a priori psychosis region of interest) and diagnoses of schizophrenia-spectrum disorder and cannabis use disorder derived from the Structured Clinical Interview for DSM-5. Linear mixed effects analyses were performed relating neuromelanin-MRI contrast to clinical measures.

**Results:** We found that CUD was associated with elevated neuromelanin-MRI signal in a cluster of ventral SN voxels (387 of 2060 SN voxels, p_corrected_=0.027, permutation test). Furthermore, CUD was associated with elevated neuromelanin-MRI signal in an SN subregion previously documented to have elevated signal in relation to untreated psychotic symptoms (t_92_ =2.12, p=0.037). In contrast, FES was not associated with a significant alteration in neuromelanin-MRI signal (241 SN voxels had elevated signal, p_corrected_=0.094).

**Conclusions and Relevance:** These findings suggest that elevated dopamine function in a critical SN subregion may contribute to the risk of psychosis in people with CUD. Thus, cannabis affects the long-suspected ‘final common pathway’ for the clinical expression of psychotic symptoms. Imaging the dopamine system with neuromelanin-MRI may index long-term dopamine turnover.

**Key Points:** *Question:* Is the same midbrain dopamine pathway impacted by cannabis as in psychosis?

*Findings:* Cannabis use disorder participants had elevated neuromelanin-MRI signal in a cluster of ventral substantia nigra voxels and in a subregion previously documented to have elevated signal in relation to untreated psychotic symptoms.

*Meaning:* Increased dopamine functioning in the ventral substantia nigra may contribute to the risk of psychosis in people with cannabis use disorders.

## Introduction

Cannabis use disorder (CUD) is a common disorder that is of particular concern due to the association between cannabis use and psychosis ^1,2^. Studies have shown that in healthy individuals, administration of Δ9-tetrahydrocannabinol (THC), the main psychoactive compound in cannabis, induces positive psychotic symptoms, such as suspiciousness, delusions, and altered perception ^3–5^ and negative symptoms, including blunted affect and amotivation ^6^. Additionally, higher levels of cannabis use are consistently associated with an increased risk of psychosis demonstrating a dose-response relationship, supporting a causal influence ^6^ and individuals with psychosis who use cannabis have an earlier illness onset compared to those who do not ^7^. However, the mechanism by which cannabis brings about this effect is still obscure and cannabis use is neither a sufficient nor necessary cause for persistent psychotic disorders such as schizophrenia ^8^. One possibility is that cannabis affects the same ‘final common pathway’ of dopaminergic excess in psychosis ^9,10^. Importantly, given its association with psychosis relapses in patients with schizophrenia ^11^, its effect on dopamine may be dose-dependent and sustained when use is continued.

The dopamine hypothesis of schizophrenia states that striatal hyperdopaminergia underlies the positive psychotic symptoms of schizophrenia ^12^. The effect of cannabis use on the dopaminergic system is less well-known. Following THC administration, dopamine is released in striatal and cortical brain areas ^13,14^; an effect similar to dopamine levels reported in psychosis and schizophrenia ^15^. In contrast, positron emission tomography (PET) studies in people with cannabis use have found reduced dopamine synthesis capacity in the striatum ^16^. Therefore, no consistent dopamine availability changes have been demonstrated in PET studies of cannabis users to date. The reported differences in short-versus long-term effects of cannabis use on dopamine further complicate the mechanistic link connecting cannabis, dopamine, and schizophrenia.

Neuromelanin-sensitive magnetic resonance imaging (neuromelanin-MRI) is an approach with non-invasive, short acquisition periods that provides an indirect index of dopamine from the substantia nigra (SN, including the ventral tegmental area (VTA)) where most dopaminergic cells originate. Neuromelanin, a breakdown product of cytosolic dopamine, accumulates in these neurons by forming insoluble complexes with iron ^19^. The resulting paramagnetic properties create an endogenous localised contrast in the MRI, quantified as contrast-noise-ratio (CNR) in comparison to a non-dopaminergic reference region. Across the lifespan, neuromelanin accumulation leads to an age-related increase of the neuromelanin-MRI signal, before dropping with the onset of neurodegeneration ^19,21,22^. Higher neuromelanin-MRI signals occur in disorders where hyperdopaminergia is suspected, such as schizophrenia ^23^ and cocaine use disorder ^24^. In schizophrenia, a subset of voxels in the SN has been identified to show higher neuromelanin-MRI CNR that correlates with psychosis severity ^23^. Thus, neuromelanin-MRI of the SN provides a viable method for examining the putative dopaminergic effects of cannabis use in relation to schizophrenia. To our knowledge, there are no studies examining the effect of CUD on neuromelanin-MRI in the SN or the SN regions in psychosis where dopaminergic turnover is likely.

We aim to investigate if neuromelanin is altered in those with CUD compared to those without CUD (nCUD), and if CUD and first episode schizophrenia (FES) have an additive effect of increasing neuromelanin. We also examined the longitudinal effect of CUD on neuromelanin over 1-year. We hypothesize that: (1) individuals with CUD will differ from nCUD in the SN neuromelanin-MRI signal, especially in regions previously shown to be related to psychosis severity; (2) neuromelanin-MRI signal abnormalities will relate to psychotic symptom burden; and (3) neuromelanin-MRI signal will reduce over 1-year in individuals with CUD compared to nCUD.

## Methods

### Participants

This study was approved by Western University’s Health Science Research Ethics Board. 18-35 years old participants were recruited from London, Ontario: Patients with FES (CUD and nCUD) from a specialized Early Intervention Program for Psychosis (PEPP-London) and community-living non-clinical volunteers (CUD and nCUD) from the same locality. The Structured Clinical Interview for DSM Disorders (SCID) was used by a research psychiatrist to diagnose schizophrenia and CUD, and to rule out current alcohol and stimulant drug use disorder. Non-clinical volunteers with CUD,were excluded if they had a first-degree family history of schizophrenia or bipolar illness. Twenty-five CUD participants and 36 nCUD participated in the study, 28 of whom had FES (Table 1; more details in Online Materials).

**Table 1:**
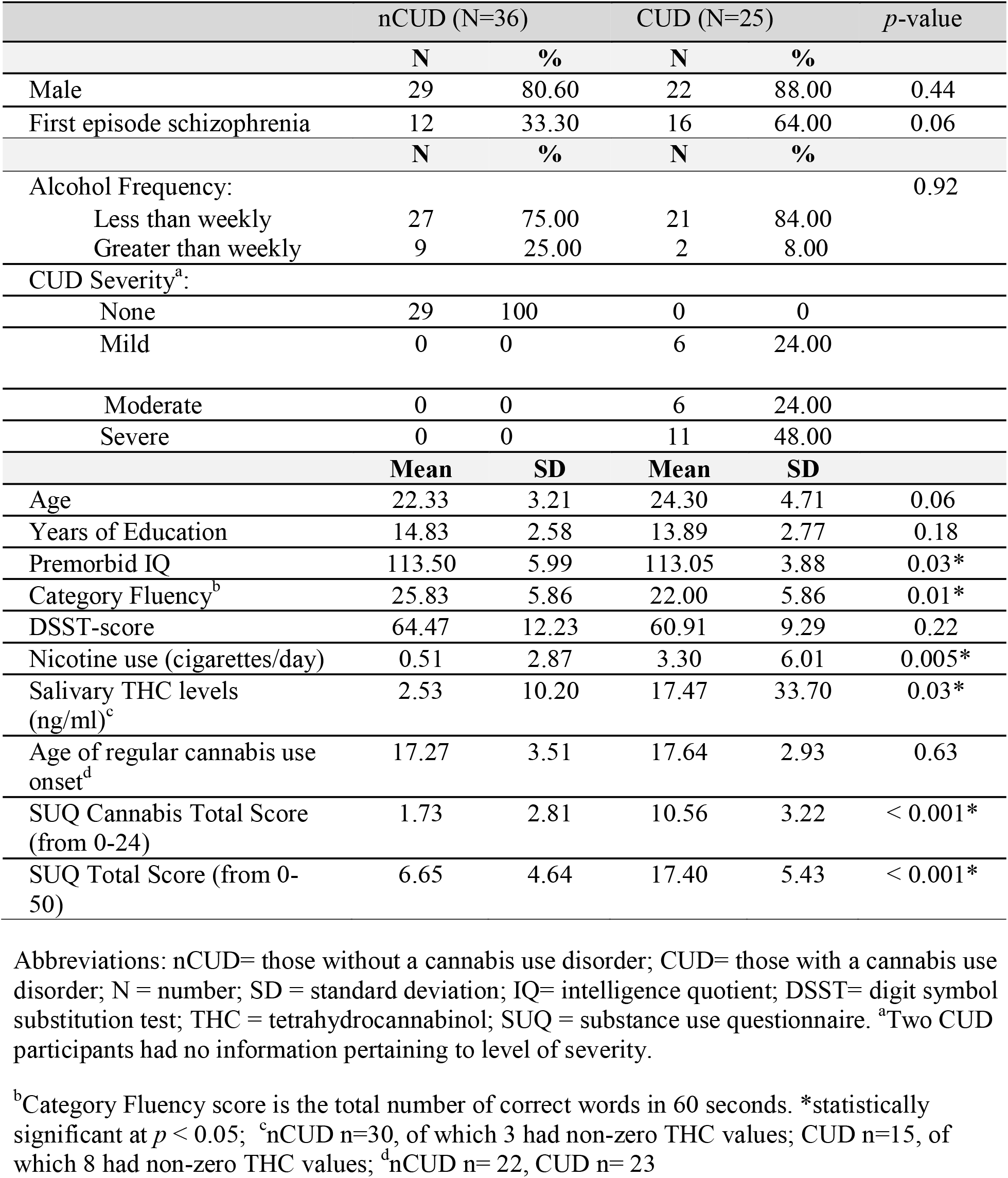
Demographics and clinical characteristics of participants.

### Clinical and Cognitive Measures

Psychotic symptoms were assessed using the Positive and Negative Syndrome Scale (PANSS; ^25^) (Table 1) for all participants, followed by cognitive assessments with National Adult Reading Test (NART), the written and oral digit symbol substitution test (W-DSST and O-DSST, mean reported together as modified-DSST; ^28^) and the Category Fluency Test (Table 1) at baseline, 6 months, and 12-month follow-up. THC levels were quantified on the day of scanning with the passive drool method (Quantisal™ device from Immunalysis) using commercially available liquid chromatography-mass spectrometry (East Coast Mobile Medical Inc.). Details of questionnaire-based substance use assessments can be found in Online Materials.

### MRI Acquisition

Magnetic resonance images were acquired for all participants on a 3T GE Discovery MR750 using a 32-channel, phased-array head coil. Neuromelanin-MRI images were collected via a two-dimensional gradient echo sequence with the following parameters: repetition time (TR) = 285 msec; echo time (TE) = 3.9 msec; flip angle = 40°; field of view (FoV) = 220 × 165 mm^2^; number of slices = 10; slice thickness = 3.0 mm; number of averages = 8; acquisition time = 12.16 minutes. The image stack was oriented along the anterior-commissure-posterior-commissure line, providing coverage of the SN-containing portions of the midbrain (and structures surrounding the brainstem) with high in-plane spatial resolution. Whole-brain, high-resolution structural MRI scans were acquired for preprocessing of the neuromelanin-MRI data: a T1-weighted 3D spoiled gradient recall echo sequence (inversion time = 400 ms; TR = 6.7 msec; TE ∼ 3.0 ms; flip angle = 11°; FoV = 256 × 256 mm^2^; matrix = 256 × 256; number of slices = 184; isotropic voxel size = 1.0 mm^3^) and a T2-weighted CUBE sequence (TR = 2500 ms; TE = 60 ms; echo train length = 100; FoV = 256 × 256 mm^2^; number of slices = 184; isotropic voxel size = 1.0 mm^3^). Neuromelanin-MRI images were visually inspected for artifacts immediately upon acquisition with scans repeated when needed.

### Neuromelanin-MRI Analysis

For neuromelanin-MRI preprocessing steps, see Supplementary Materials. All analyses were carried out in Matlab (Mathworks) using custom scripts. Robust linear regression analyses were performed across subjects for every voxel *v* within the SN mask, as: *CNR*_*V*_ = *β*_0_ + *β*_1_ *measure of interest* + 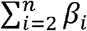 *nuisance covariate* + *ε*. Nuisance covariates, including diagnosis, and age, varied for different analyses; while all analyses included age, diagnosis (FES or CUD) was only included as a covariate in analyses where these variables differed across subjects. Robust linear regressions were used to minimize the need for regression diagnostics in the context of mass-univariate, voxelwise analyses. Voxelwise analyses were carried out within the template SN mask after censoring subject data points with missing values or extreme values [values more extreme than the first or the 99th percentile of the CNR distribution across all SN voxels and subjects (CNR values below −9% or above 40%)]. In the voxelwise analyses, the spatial extent of an effect was defined as the number of voxels *k* (adjacent or nonadjacent) exhibiting a significant relationship between the measure of interest and CNR (voxel-level height threshold for t-test of regression coefficient β1 of *p* < 0.05, one-sided [β∗1]). Hypothesis testing was based on a permutation test in which the measure of interest was randomly shuffled with respect to CNR. This test corrected for multiple comparisons by determining whether an effect’s spatial extent *k* was greater than would be expected by chance (*p*_corrected_ < 0.05; 10,000 permutations; equivalent to a cluster-level familywise error-corrected *p* value). On each iteration, the order of the values of a variable of interest (e.g., CUD status) was randomly permuted across subjects (and maintained for the analysis of every voxel within the SN mask for a given iteration of the permutation test, accounting for spatial dependencies). This provided a measure of spatial extent for each of the 10,000 permuted datasets, forming a null distribution against which to calculate the probability of observing the spatial extent *k* of the effect in the true data by chance (*p*_corrected_).

As we can make a more direct link between cannabis use and psychotic experiences than with the diagnosis of schizophrenia per se ^29^, and to remove the risk of circularity when subregional selection and data analysis are performed on the same dataset, we focused on the neuromelanin-MRI signal of the voxels correlated with psychosis symptom severity in Cassidy et al. ^23^ (‘psychosis voxels’) for a Region-of-Interest analysis and to study the dose effect.

Given the known association of cannabis use to negative symptoms and general psychopathology ^29^, we assessed if these symptoms vary with the mean neuromelanin-MRI CNR extracted from the ‘CUD voxels’, where the most prominent increase in neuromelanin signal occurred in the voxelwise search in our sample.

## Results

Baseline demographic and clinical characteristic data is shown in Table 1. In total, 25 individuals with a CUD and 36 nCUD individuals were included. No significant group differences were found for sex, age, or years of education (*p* > 0.05) (Table 1). The CUD group had significantly lower premorbid IQ scores (*p*= 0.03) and category fluency scores (*p*= 0.01) than the nCUD group (Table 1). Those with a CUD had significantly increased nicotine use (*p* = 0.005) and salivary THC levels (*p* = 0.01) compared to the nCUD group (Table 1). No significant differences were found between alcohol use frequency, age of regular cannabis use onset, or modified-DSST scores (*p* > 0.05). (Table 1). There were more FES participants in the CUD group than nCUD, with trend-level significance (Table 1). Demographic/clinical characteristics for all groups can be found in eTable 1.

### Voxelwise association between neuromelanin-MRI signal, CUD, and psychosis

In a voxelwise analysis, CNR was higher in CUD participants compared to nCUD participants in a ventral SN voxel cluster (387 of 2060 SN voxels at *p*<0.05, linear mixed-effects analysis controlling for FES diagnosis, age, and sex, p_corrected_=0.027, permutation test; Figure 1 ‘CUD voxels’). There was also a cluster in the mediodorsal SN where neuromelanin-MRI signal was lower in CUD participants, although this cluster did not achieve significance (211 of 2060 SN voxels, p_corrected_=0.15, permutation test; Figure 1). On the other hand, in this model, FES diagnosis was not significantly associated with neuromelanin-MRI signal (241 of 2060 SN voxels showed increased signal in FES participants, p_corrected_=0.094, permutation test). Furthermore, there was no significant evidence of an interaction of CUD by FES (129 of 2060 SN voxels showed a positive interaction, p_corrected_=0.30; 62 of 2060 SN voxels showed a negative interaction, p_corrected=_ 0.47). There was no effect of time on SN signal in any groups. For details, see Online Materials.

**Figure 1.**
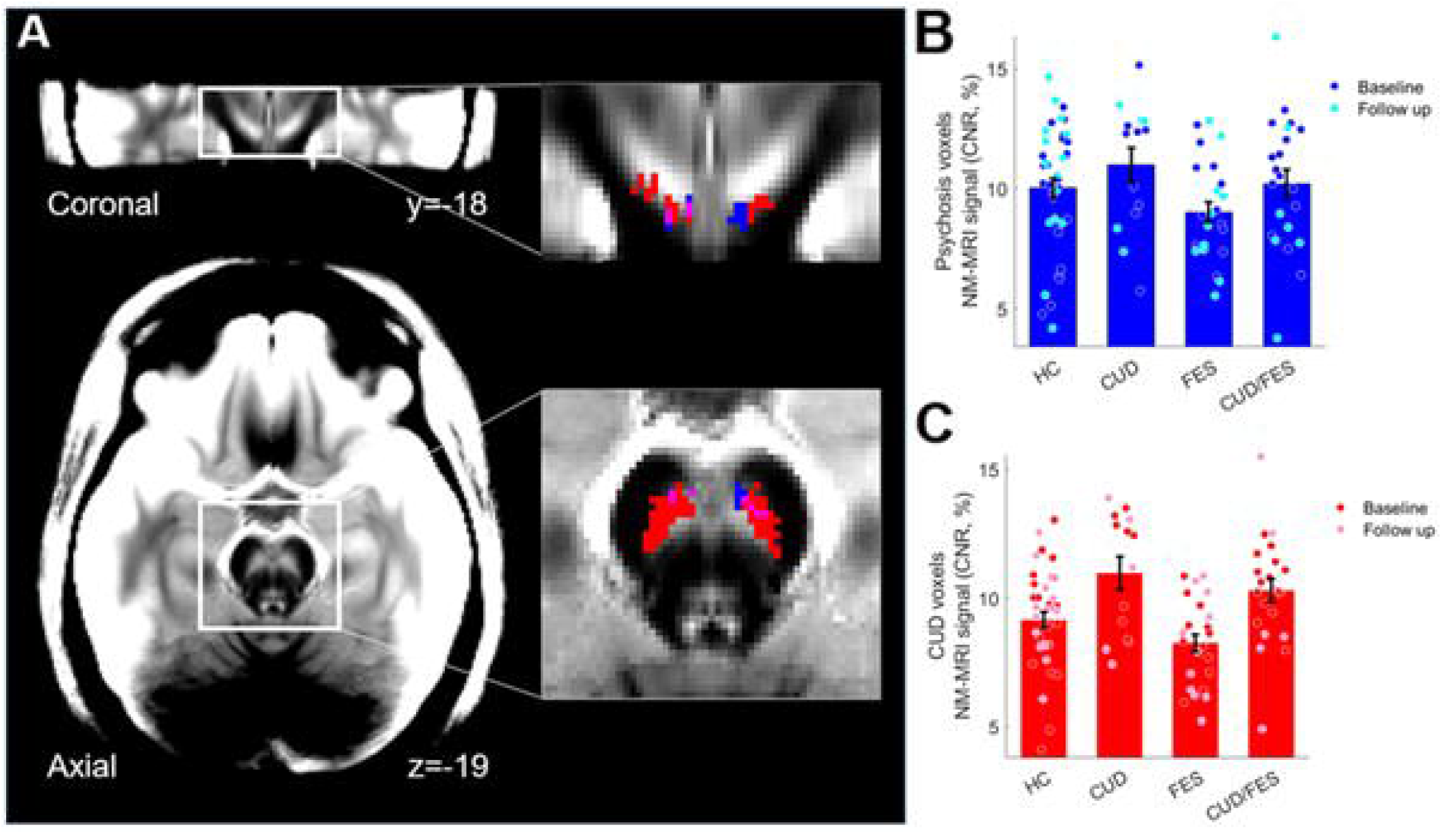
Substantia nigra neuromelanin-sensitive magnetic resonance imaging (NM-MRI) signal in cannabis use disorder and first episode schizophrenia participants. (A) A visualization template of the midbrain generated by averaging spatially normalized neuromelanin-MRI images from study participants. Magnifications show substantia nigra voxels where neuromelanin-MRI signal was elevated (red) in cannabis use disorder (CUD) participants compared to individuals without a cannabis use disorder (nCUD). Psychosis voxels previously shown to correlate with untreated positive symptoms of psychosis are shown in blue (overlap with cannabis-elevated voxels in violet). These voxels were clustered near the cannabis-elevated voxels. (B) Dot-plots showing neuromleanin-MRI signal extracted from psychosis voxels in four groups, including healthy controls (HC), those with a CUD, those with first episode schizophrenia (FES), and those with both a CUD and FES. Blue dots represent scans collected at baseline and cyan dots at follow-up. (C) Dot-plots showing neuromelanin-MRI signal extracted from cannabis voxels in four groups; red dots were collected at baseline and pink dots at follow-up. Error bars represent standard error of the mean.

### Region of interest analysis

Neuromelanin-MRI signal extracted from a mask of ‘psychosis voxels’ was found to be significantly higher in CUD participants compared to nCUD (t_92_=2.12, p=0.037, linear mixed effects model controlling for FES diagnosis, age, and sex; Figure 1). There was no interaction between FES diagnosis and CUD in these voxels (t_55_=0.02), though the CUD effect on the neuromelanin signal was numerically stronger in those with FES (Cohen’s d= 0.71) compared to those without FES (Cohen’s d=0.39; Figure 1).

To test if increasing CUD severity linearly relates to a change in CNR within the ‘psychosis voxels’, we conducted a regression analysis with CUD severity grouped as none, mild, or moderate/severe. This grouping, though arbitrary, provided the optimal distribution to test the dose effect. In the ‘psychosis voxels’, we found a significant dose-dependent relationship between neuromelanin CNR and CUD (F(1, 96) = 4.89, p = 0.029; Figure 2). This indicates that an escalating severity of CUD may contribute to increased neuromelanin signal in midbrain regions that are most sensitive to psychotic symptom burden.

**Figure 2.**
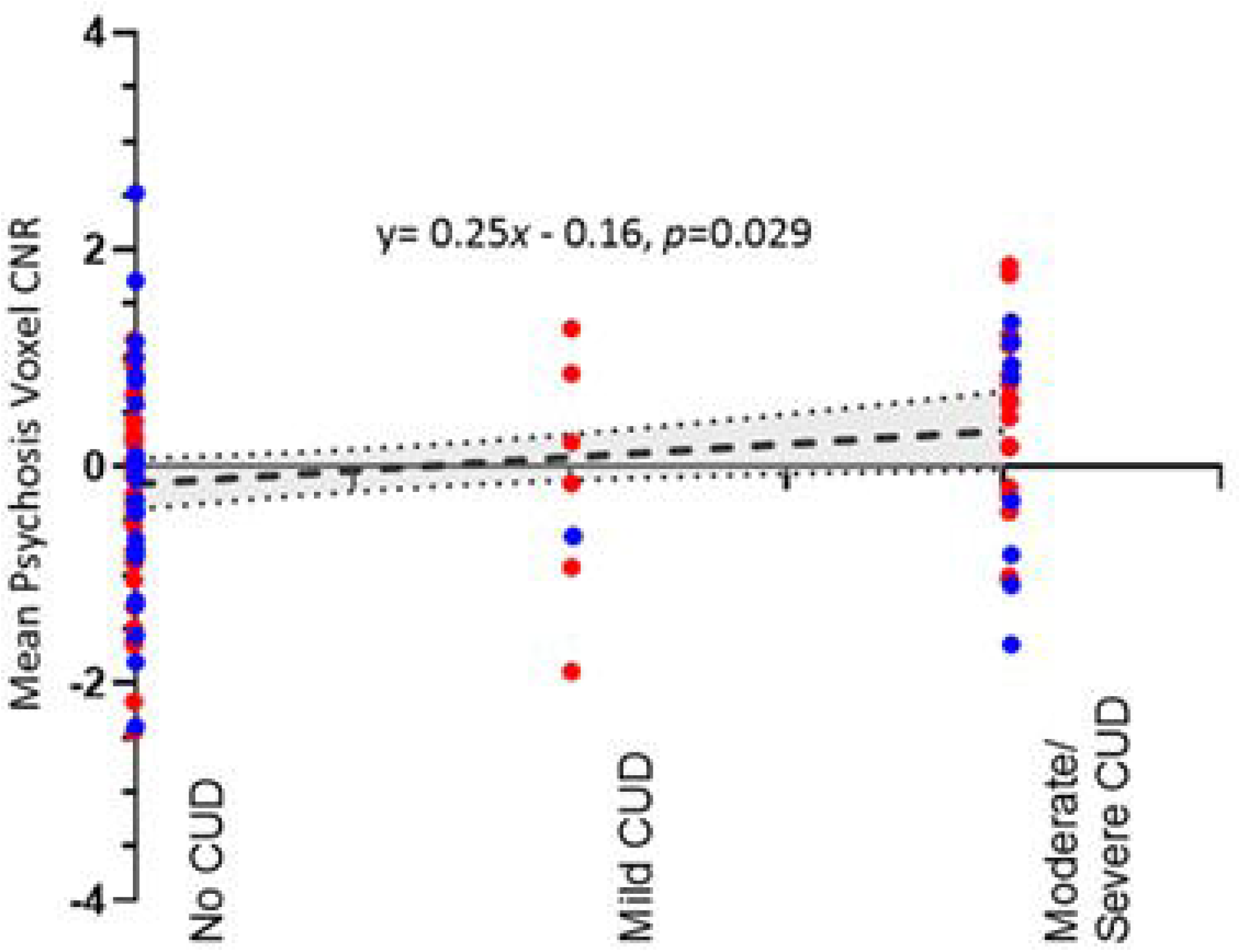
Psychosis voxel signal and cannabis use disorder severity. Correlation between mean neuromelanin-MRI contrast-to-noise ratio in ‘psychosis voxels’, corrected for age, sex, and diagnosis, and severity of cannabis use disorder.

## Discussion

We examined the neuromelanin-MRI signal in those with a CUD and those with FES in a voxelwise manner. We located a ventral SN region where CUD had an independent effect on higher neuromelanin. Individuals with CUD also had higher neuromelanin in the subset of voxels with prior association with psychosis severity ^23^. Based on data demonstrating SN neuromelanin accumulation is dependent on dopaminergic functioning ^23,30^, our results suggest that elevated neuromelanin CNR signal in SN subregions of individuals with a CUD may be an indication of dopamine dysfunction. This dopaminergic dysfunction occurs within the same area of SN where higher neuromelanin accumulation occurs in people with more severe symptoms of psychosis.

Thus, cannabis use may exacerbate the dopaminergic dysfunction that is relevant for psychosis. This is consistent with 3 lines of evidence: (1) clinical studies that link cannabis use with more pronounced positive symptoms in those with psychosis ^8^. (2) preclinical research indicating striatal dopamine receptor upregulation with THC use ^31^, demonstrating nigrostriatal pathway dysfunctions with cannabis use; and (3) imaging studies in schizophrenia showing increased dopamine in the dorsal striatum, which receives SN projections of the nigrostriatal pathway ^14^, consistent with having increased dopamine in the SN and thus assumed to correlate with increased neuromelanin as demonstrated here. Our results suggest that these changes are related, providing further insight into the neurochemistry of cannabis use with psychosis. While we did not see a notable increase in neuromelanin-MRI in those with FES compared to HCs, likely due to a lack of power, we did see an increase in voxels similar to those found previously (eFigure 2)^32^.

Alongside the higher neuromelanin CNR in ‘psychosis voxels’ of individuals with CUD, we identified a notable increase in neuromelanin CNR in ventral SN voxels. Despite the literature suggesting decreased striatal dopaminergic activity with long-term cannabis use, many variables likely tie into the relationship between dopamine and cannabis use. Many PET studies have found no differences in striatal dopamine receptor availability with cannabis use ^18,33–35^. However, these studies were associated with relatively low dependence severity rather than assessing severe cannabis dependence. CUD participants in our study had an average SUQ cannabis score of 10.56 (SD=3.22) out of the possible 24, suggesting most individuals had a moderate dependence. Additionally, average age of cannabis use onset in those with a CUD was 17.64 (SD=2.93), whereas average age at baseline was 24.32 (SD= 4.78); thus, on average, it had been less than 10 years since cannabis use onset. We do not see a significant CUD-related reduction in midbrain neuromelanin signal in our sample.

It has been previously reported that neuromelanin levels within the SN increase with age ^36–38^. The age range in our study was small, ranging from 17 to 35 (mean 22.78 ± 3.96), thus reducing any age-related changes. This may explain our null findings in relation to neuromelanin CNR after the 1-year period. Additionally, based on the acute dopamine increase as well as reduced dopamine release in more established users seen in PET studies, we expected neuromelanin CNR to decrease over 1-year in CUD compared to nCUD. But, we did not see any effect of time by CUD, nor any interaction between CUD, psychosis, and time. This could be attributed to the fact that individuals in the CUD group continued to satisfy persistent CUD at both time points, with no attrition of diagnosis. Consequently, it is plausible that they had already undergone the blunting effects of CUD prior to their initial evaluation, thereby precluding further significant decline in dopamine blunting.

While neuromelanin has been mostly seen as a product of dopamine auto-oxidation, it is not completely inert ^39^. It binds to metals and toxic compounds that contribute to oxidation-reduction processes, and may prevent neuronal damage, partly as an antioxidant and scavenger of radical ions ^39^. In addition, neuromelanin can function as an iron-binding molecule to regulate iron homeostasis in neuromelanin-containing neurons ^40^. Thus, we cannot infer if higher or lower levels of neuromelanin represent ill versus beneficial effects. We express caution in making such inferences especially in relation to the effects of cannabis and psychosis on the neuromelanin signal.

Some methodological decisions we made in this work include: (1) analyzing a subset of voxels based on a previous neuromelanin study ^23^; and (2) examining mean neuromelanin CNR for specific SN subsets. The use of an external source for the voxel subset was chosen to prevent circular reasoning, such that we ensure the evidence for psychosis is independent of our dataset, and because it is based upon unmedicated participants, whereas our dataset includes individuals with antipsychotic use, which may impact neuromelanin CNR. However, we recognize that this decision may introduce its own limitations, including that the relationship to symptom severity of these voxels have not been reproduced. Secondly, the use of voxel subsets instead of a mean value for the entire SN was decided because distinct areas of the SN project to distinct regions of the nigrostriatal system ^41^; thus, the use of an overall mean may not be representative of a useful metric.

Our study has several noteworthy strengths. First, we employed means of voxel groups and voxelwise analyses of neuromelanin CNR, ensuring a comprehensive evaluation of its distribution and potential abnormalities. Additionally, the robustness of our study is reinforced by the many methods of examining cannabis use; CUDs were diagnosed by a psychiatrist, and we included both salivary THC and scores from the cannabis subscale of the SUQ, which enhances the reliability and accuracy of the findings. Despite these strengths, certain limitations exist. One notable limitation is the underrepresentation of female participants, affecting the generalizability of our results. Furthermore, we focused exclusively on the SN/VTA region and did not explore the locus coeruleus, another neuromelanin-rich region. Our subsample with longitudinal data was powered only to detect large effect changes over time (effect size of 0.8 or greater); thus, the lack of change over the 1-year period needs to be treated with caution. Addressing these limitations in future research would contribute to a more comprehensive understanding of how schizophrenia and CUD affect dopamine turnover.

CUD is associated with increased neuromelanin in voxels that are relevant for the severity of psychosis, providing a potential mechanistic lead for the observed effects of cannabis on psychotic symptoms. The presence of higher neuromelanin in certain voxels of the SN in FES and lower neuromelanin in certain voxels in CUD requires further validation in larger studies and with longitudinal data.

## Acknowledgements

The following individuals provided important contributions, without which this work would not have been possible: Kyle McKee, Candice Crocker, Sherry Stewart, James Rioux, Paulina Dzialozynski. J Ahrens reports funding from the Canadian Institutes of Health Research, Schizophrenia Society of Canada Foundation and the Canadian Consortium for Early Intervention in Psychosis, the Fonds de Recherche du Quebec—Santé and Quebec Bio-Imaging Network (QBIN). R A Rabin received funding from the Fonds de Recherche du Quebec—Santé. L Palaniyappan’s research is supported by the Canada First Research Excellence Fund, awarded to the Healthy Brains, Healthy Lives initiative at McGill University (through New Investigator Supplement) and Monique H. Bourgeois Chair in Developmental Disorders. He receives a salary award from the Fonds de recherche du Quebec-Sante. A R Khan’s research is supported by the Canada First Research Excellence Fund, awarded to the BrainsCAN program at Western University, the Canada Research Chairs program #950-231964, and Canada Foundation for Innovation (CFI) John R. Evans Leaders Fund project #37427. Data acquired for this study was funded by the Canadian Institutes of Health Research Project Grant (371730) to P Tibbo, with L Palaniyappan, K McKee; CE Crocker; AR Khan and SH Stewart. LP reports personal fees from Janssen Canada, Otsuka Canada, SPMM Course Limited, UK, Canadian Psychiatric Association; book royalties from Oxford University Press; investigator-initiated educational grants from Sunovion, Janssen Canada, Otsuka Canada outside the submitted work. P Tibbo has honoraria for advisory boards and speaker fees for Janssen, Otsuka Lundbeck and investigator-initiated reach grant from Janssen. CMC reports an investigator-initiated sponsored research agreement from Terran Biosciences outside the submitted work and having filed patents for the analysis and use of neuromelanin-sensitive MRI in central nervous systems disorders licensed to Terran Biosciences with no royalties received.

