## Supplemental Materials for "Converging effects of cannabis and psychosis on the dopamine system: A longitudinal neuromelanin-sensitive MRI study in cannabis use disorder and first episode schizophrenia"

### **Online Materials**

eTable 1. Demographics and clinical characteristics of the cannabis use disorder group and non-cannabis use disorder group, with and without first episode schizophrenia.

eFigure 1. Mean neuromelanin-MRI contrast-to-noise ratio, corrected with age, sex, and psychosis diagnosis, in the substantia nigra subregion increased in cannabis use disorder and the relationship with PANSS negative scores. Red dots are baseline values and blue dots are follow-up values.

eFigure 2. Substantia nigra voxels in which neuromelanin-MRI signal was elevated in first episode schizophrenia relative to controls.

### **Supplementary Methods**

**Participants**

The diagnostic assessment of the SCID was repeated at 6 months and 12 months later to confirm the diagnosis of schizophrenia and CUD. Timeline Followback method was used to collect detailed information about past and current use of cannabis in all subjects at baseline, 6 months and 1 year. The two groups were stable over the 1 year, with all CUD subjects still satisfying the CUD criteria and no new diagnosis of CUD emerging in the non-CUD group in this sample. CUD participants were grouped to have a mild, moderate, or severe CUD based upon the DSM-5 criteria for substance use disorders, specific to cannabis. If participants met two or three of the 11 DSM-5 symptoms, they were indicated to have a mild CUD, four or five as moderate, and six or more symptoms as severe ^1^. Individuals with known progressive brain diseases (e.g., demyelinating), contraindications to MRI, significant head injury, seizures, or ongoing alcohol or stimulant drug use disorder were excluded.

**Clinical and Cognitive Measures**

Premorbid IQ was measured with the National Adult Reading Test (NART; ^2^). Alcohol use, cannabis use, nicotine use, and drug dependency were measured with the Alcohol Use Disorders Identification Test (AUDIT; ^3^), Cannabis Abuse Screening Test (CAST; ^4^), Fagerstrom Test for Nicotine Dependence (FTND; ^5^), and the Substance Use Questionnaire (SUQ; ^6^).

**Neuromelanin-MRI Preprocessing**

Neuromelanin-MRI scans were preprocessed using SPM12 software to allow for voxelwise analyses in standardized Montreal Neurological Institute (MNI) space. Neuromelanin-MRI scans were coregistered to participants’ T1-weighted scans and tissue segmentation was performed using the T1-weighted images. Neuromelanin-MRI scans were normalized to MNI space using DARTEL routines with a gray- and white-matter template generated from all study participants. The resampled voxel size of unsmoothed, normalized neuromelanin-MRI scans was 1 mm, isotropic. Intensity normalization and spatial smoothing with a 1 mm full-width-at-half-maximum Gaussian kernel were performed using custom MATLAB (MathWorks) scripts. Neuromelanin contrast-to-noise ratio (CNR) for each of the 2060 SN voxels (*v*) in each participant was calculated as change in neuromelanin-MRI signal intensity (*I*) to a reference region (*RR*), the crus cerebri, white-matter tracts known to have minimal neuromelanin: CNR*_v_*=(*I_v_* – mode(*I_RR_*))/mode(*I_RR_*).

A template mask of the reference region and of the SN was created by manual tracing on a template neuromelanin-MRI image in MNI space (an average of normalized NM-MRI scans from all study participants). The mode*(I_RR_)* was calculated for each participant from a kernel-smoothing function fitted to a histogram of the distribution of all voxels in the mask. The resulting neuromelanin-MRI contrast-to-noise ratio maps were then spatially smoothed with a 1-mm full width at half maximum Gaussian kernel.

**Supplementary Results**

**eTable 1. Demographics and clinical characteristics of the cannabis use disorder group and non-cannabis use disorder group, with and without first episode schizophrenia.**

|  |  | nCUD-FES  (N=12) | | CUD-FES  (N=16) | | nCUD-nFES  (N=24) | | CUD-nFES  (N=9) | | *p*-value |
| --- | --- | --- | --- | --- | --- | --- | --- | --- | --- | --- |
|  |  | **N** | **%** | **N** | **%** | **N** | **%** | **N** | **%** |  |
|  | Male | 10 | 83 | 14 | 88 | 19 | 79 | 8 | 89 | 0.89 |
|  | Alcohol Frequency: |  |  |  |  |  |  |  |  | 0.76 |
|  | | nCUD-FES  (N=12) | | CUD-FES  (N=16) | | nCUD-nFES  (N=24) | | CUD-nFES  (N=9) | | *p*-value |
|  | | **N** | **%** | **N** | **%** | **N** | **%** | **N** | **%** |  |
|  | Never | 1 | 8 | 4 | 25 | 3 | 12.5 | 0 | 0 |  |
|  | Monthly | 5 | 41.7 | 7 | 44 | 9 | 37.5 | 4 | 44 |  |
|  | 2-4 times per month | 3 | 25 | 3 | 19 | 5 | 20.8 | 3 | 33 |  |
|  | 2-3 times per week | 2 | 16.7 | 1 | 6.3 | 5 | 20.8 | 2 | 22 |  |
|  | >4 times per week | 0 | 0 | 0 | 0 | 2 | 8.3 | 0 | 0 |  |
|  |  | **Mean** | **SD** | **Mean** | **SD** | **Mean** | **SD** | **Mean** | **SD** |  |
|  | Age | 23.0 | 2.17 | 24 | 5.1 | 22.0 | 3.61 | 24.33 | 3.74 | 0.43 |
|  | Years of Education | 13.33 | 1.73 | 12.8 | 1.96 | 15.40 | 2.65 | 16.81 | 2.59 | 0.0002* |
|  | Premorbid IQ | 108.85 | 3.67 | 107.38 | 6.29 | 115.52 | 5.72 | 113.05 | 3.87 | 0.0001 |
|  | Nicotine use (cigarettes/day) | 5.92 | 9.0 | 5.0 | 7.42 | 0 | 0 | 0 | 0 | 0.003* |
|  | Category Fluency score** | 21.33 | 8.14 | 16.8 | 5.26 | 25.83 | 5.86 | 22.0 | 5.86 | 0.0007* |
|  | DSST score | 45.42 | 9.76 | 44.47 | 8.80 | 65.15 | 12.20 | 61.18 | 10.86 | <0.0001* |
|  | GAF score | 54.25 | 17.82 | 52.29 | 12.32 |  |  |  |  | 0.67 |
|  | PANSS positive | 10.66 | 3.58 | 13.73 | 4.45 |  |  |  |  | 0.07 |
|  | PANSS negative | 11.42 | 3.29 | 13.2 | 5.16 |  |  |  |  | 0.33 |
|  | PANSS general | 25.92 | 6.69 | 26.07 | 7.09 |  |  |  |  | 0.98 |
|  | SOFAS score | 53.83 | 15.66 | 49.93 | 18.24 |  |  |  |  | 0.60 |
|  | CGI-S score | 4.36 | 1.63 | 4.29 | 1.03 |  |  |  |  | 0.92 |
|  | Days since illness onset*** | 110.6 | 59.38 | 87.88 | 52.11 |  |  |  |  | 0.36 |
|  | Rx adherence % | 93.75 | 21.65 | 90.63 | 18.42 |  |  |  |  | 0.63 |
|  | DDD | 1.08 | 0.70 | 1.03 | 0.76 |  |  |  |  | 0.87 |
|  | Salivary THC levels (ng/ml)**** | 5.92 | 9.0 | 16.82 | 32.09 | 0.32 | 1.38 | 8.5 | 9.8 | 0.11 |
|  | Age of regular cannabis use onset***** | 17.86 | 2.67 | 17.12 | 3.15 | 17.69 | 3.71 | 18.56 | 2.79 | 0.66 |

Abbreviations: PT= first episode schizophrenia patients; HC= healthy controls; nCUD= without a cannabis use disorder; CUD= with a cannabis use disorder; N= number; SD= standard deviation; IQ= intelligence quotient; DSST= digit symbol substitution test; GAF= global functioning scale; PANSS= positive and negative syndrome scale; SOFAS= social and occupational functioning scale; CGI-S= clinical global impressions-severity; Rx= prescription; DDD= defined daily dose (of antipsychotic medications); THC = tetrahydrocannabinol.

**p* <0.05 **raw count ***equal to days between admission date to psychosis program and the date of the study visit ****PT-nCUD n=7, PT-CUD n=11 , HC-nCUD n=19, HC-CUDn=4. *****in years, PT-nCUD n=7, PT-CUD n=15, HC-nCUD n=13, HC-CUD n=10.

*Voxelwise association between neuromelanin-MRI signal, CUD, and psychosis*

There was no effect of time on SN signal (46 of 2060 voxels decreased over time, p_corrected_=0.82, permutation test; follow-up N=37), nor a significant time by CUD [86 of 2060 voxels decreased, p_corrected_=0.52; 55 of 2060 increased, p_corrected_=0.76; nCUD n=25; CUD= 12], or time by FES interaction [138 voxels decreased over time, p_corrected_=0.41; 27 voxels increased over time, p_corrected_=0.83; HC n= 19; FES n= 18], or interaction between CUD, psychosis diagnosis, and time [118 voxels decreased over time, p_corrected_=0.38, 48 voxels increased over time, p_corrected_=0.76].

*Association between CUD-related neuromelanin-MRI signal and symptom burden*

The mixed model analysis revealed a trend between mean neuromelanin CNR from ‘CUD voxels’ and PANSS negative scores, suggesting that with each point incremental change in PANSS negative score, the projected value of mean neuromelanin CNR escalates by 0.136 units (under the condition that all other variables remain unchanged). This observed trend, though not achieving conventional significance (F(1,93) = 0.136, p = 0.07; with age and CUD status as covariates), hints at a potential association worthy of further exploration (see Figure S1). No such relationships were noted between neuromelanin CNR of the ‘CUD voxels’ and PANSS positive (F(1,84) = 0.61, p > 0.05) or general (F(1,84) = 0.25, p > 0.05) symptoms. No relationships were found between neuromelanin CNR extracted from the ‘psychosis voxels’ and PANSS negative (F(1,96) = 0.43, p > 0.05), positive (F(1,84) = 0.15, p > 0.05), or general (F(1,84) = 0.1, p > 0.05) symptoms. Taken together, the trend reported here calls for further investigations to clarify if the CUD-related increase in neuromelanin signal contributes to a higher negative symptom burden, and whether this relationship is restricted to the midbrain regions that are most sensitive to cannabis use.

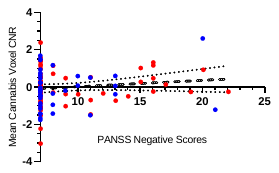

**eFigure 1. Cannabis voxel signal and PANSS negative scores.** Mean neuromelanin-MRI contrast-to-noise ratio, corrected with age, sex, and psychosis diagnosis, in the substantia nigra subregion increased in cannabis use disorder and the relationship with PANSS negative scores. Red dots are baseline values and blue dots are follow-up values. *Abbreviations:* CNR, contrast-to-noise ratio; PANSS, positive and negative syndrome scale.

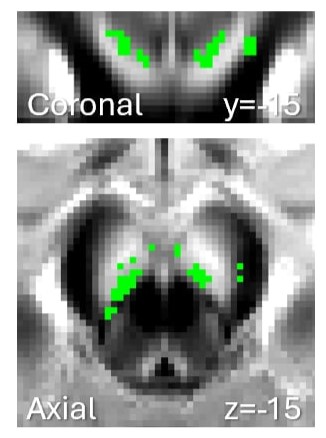

**eFigure 2.** **Substantia nigra voxels in which neuromelanin-MRI signal was elevated in first episode schizophrenia relative to controls.**
